# Borrelia Burgdorferi binds Serum Amyloid A and Modulates Subcutaneous Adipose Tissue Immune Signaling

**DOI:** 10.64898/2026.01.29.702514

**Authors:** Allen T. Esterly, Shanshan Du, Bill Lee, Emily Johnson, Anahi Odell, Esen Sefik, Qian Yu, Yingjun Cui, Haripriya Ayyala, Siba Haykal, Ian Odell, Richard Flavell, Erol Fikrig, Heidi J. Zapata

**Affiliations:** Section of Infectious Diseases, Department of Internal Medicine, Yale School of Medicine; Department of Epidemiology and Microbial Diseases, Yale School of Public Health; Department of Dermatology, Yale School of Medicine; Department of Immunobiology, Yale School of Medicine; Division of Plastic and Reconstructive Surgery, Department of Surgery, Yale School of Medicine

## Abstract

*Borrelia burgdorferi* causes over 470,000 infections annually [1]. Following a tick bite, *Borrelia* spirochetes replicate in human skin and disseminate through tissues, including subcutaneous adipose tissue. Adipose tissue is a complex organ containing non-immune and immune cells that play a significant role in the immune response [2], yet little is known about adipose tissue signaling after *Borrelia* infection. We investigated the landscape of immune signaling within adipose tissue-resident cells during *Borrelia* infection in human tissue (skin and adipose tissue) *ex vivo*. Immune pathways overall were downregulated in adipose tissue during *Borrelia* infection. Despite this, adipose stem/progenitor cells exhibited increased pro-inflammatory and extracellular matrix (ECM)-related signaling (IL-6, MIF, collagen, laminin, fibronectin), positioning them as key hubs of intercellular communication during infection. Myeloid lineage cells showed cluster-specific upregulation of immune genes such as *IL1B* and *TNF*. Network analyses highlighted laminin, and strong outgoing MIF signals in all adipose-resident cell clusters. Among the identified genes modulated after *Borrelia* infection, we observed reduced expression levels of serum amyloid A in adipose tissue-resident cells, a marker typically elevated in Lyme disease patient serum during acute infection. Here, we show that human serum amyloid A1 and serum amyloid A2 directly bind *B. burgdorferi* spirochetes, using flow cytometry. Functionally, serum amyloid A2 reduces spirochete viability in vitro in a dose-dependent manner and enhances opsonization and phagocytosis by macrophages. Together, these findings show that adipose tissue is a site of active immune signaling in the setting of *Borrelia* infection and identifies SAA as a host defense factor against *Borrelia*.

## Introduction

In the United States, *Borrelia burgdorferi* sensu stricto (*Bb*) causes Lyme Disease (LD) by targeting skin, heart, and joints as sites of infection. Disease syndromes correlate with target organs exhibited by erythema migrans, Lyme carditis, and Lyme arthritis, respectively [3]. Cutaneous immune responses characterized by microarray and single cell sequencing have been reported previously, [4, 5] yet adipose tissue has not been previously explored. *Borrelia* spirochetes preferentially bind components of the extracellular matrix (ECM) and connective tissues during infection [6], which are highly abundant in adipose tissue. Subcutaneous adipose tissue underlies nearly all human skin and is within intimate proximity of initial skin spirochete infection sites [7, 8]. Finally, adipose tissue is adjacent to all target organs in *Borrelia* infection. Specifically, articular fat pads are adjacent to joints, also common sites of Lyme-induced arthritis [3]—perisynovial adipose tissue did show evidence of infection in a study of late *Borrelia* dissemination in macaques [9]. Similarly, pericardial adipose tissue is adjacent to the heart, the organ affected during Lyme carditis. Given that adipose tissue is not just an energy reservoir but an active immune organ, its position near to these sites of infection raises critical questions about its involvement in the pathogenesis of Lyme disease.

Adipose tissue is a known reservoir for other vector-borne parasites such as *Trypanosoma cruzi, T. brucei*, and *Plasmodium* [10–12]. *Trypanosoma cruzi* and *T. brucei*, which cause Chagas disease and African sleeping sickness respectively, both invade the cytoplasm of adipocytes or interstitial spaces of adipose tissue [13–15]. Tick transmitted *Rickettsia prowazekii* and *Coxiella burnetii* also invade the adipocyte cytoplasm [16, 17]. Post-mortem tissue examinations identified subcutaneous white adipose tissue as a site of *P. falciparum-*infected RBC accumulation in adolescent cerebral malarial patients [11], reproduced in murine models of cerebral malaria [10]. Although the above-mentioned microbes are transmitted by a variety of arthropods, these highlight the potential for adipose tissue to serve as a reservoir for microbes and as a niche for potential immune evasion [12]. These studies highlight the importance of investigating the role of adipose tissue in Lyme disease, a vector transmitted disease.

Therefore, we set out to evaluate the adipose tissue immune landscape in the setting of *Borrelia* infection. Adipose tissue is an intricate cellular community that is made up of both immune (i.e. macrophages, monocytes, dendritic cells) and non-immune cells (i.e. adipocyte stem/progenitor cells (ASPCs), endothelial cells) with strong evidence to show adipose-resident cells are actively involved in the immune response [7, 8, 18]. To further explore the immune landscape of subcutaneous adipose tissue, we performed single cell RNA sequencing on human skin with subcutaneous adipose tissue infected with *Borrelia ex vivo*. We observed modulation of immune pathways, including IFN-signaling, TNFA signaling, and extracellular matrix signaling. We specifically observed downregulation of serum amyloid A, an acute phase proinflammatory serum protein that can be produced extrahepatically in adipose tissue.

Serum amyloid A family proteins (SAA1 and SAA2) are acute phase reactants that are highly elevated in LD patients [19, 20]. Acute phase SAA proteins are small (12-14kDa) proteins that are synthesized in response to inflammatory cytokine signaling in hepatocytes and released into the serum where they bind lipoproteins [21, 22]. Extrahepatic production of SAA can occur in monocytes, macrophages [23, 24], adipocytes [25–28], and in a variety of epithelial cells [29, 30]. Serum concentrations of SAA increase up to 1000-fold, as high as 1mg/mL during acute infection, injury, or other inflammatory conditions [21, 22], therefore providing a clinical diagnostic marker of inflammation, as observed in the case of early Lyme disease [19, 20]. SAA can have antimicrobial properties such as opsonization of gram-negative bacteria by binding outer membrane proteins (e.g. OmpA) [31–33]. Finally, SAA is considered an intrinsically disordered protein (IDP) due to its lack of fixed secondary structure, which allows it to attain multiple conformations, and therefore has an innate ability to bind a wide range of ligands [34].

The ability of SAA to bind a variety of ligands, elevated SAA serum levels during early Lyme disease, and the capability of SAA to opsonize gram negative bacteria made SAA an attractive candidate to pursue. Additionally, SAA2 was previously found to bind *Bb* N40 spirochetes using a curated yeast display termed bacterial selection to elucidate host-microbe interactions in high throughput (BASEHIT) [35]. Therefore, we set out to validate the interaction between spirochetes and SAA in this study. Using flow cytometry, we were able to validate the binding of *Borrelia* to SAA proteins highlighting an important role for acute phase reactants in *Borrelia* infection within adipose tissue. We further investigated the function of SAA binding *Borrelia* and demonstrated that the binding of SAA to *Borrelia* led to decreased viability of spirochete cultures over time and enhanced phagocytosis *in vitro*. Taken together, our results show that SAA is an important host defense molecule against *Borrelia* infection and that inflammatory responses in subcutaneous adipose tissue play a role in *Borrelia* pathogenesis.

## Results

### Bb Infection Modulates Immune Pathways in Human Subcutaneous Adipose

To further investigate the immune response of skin and subcutaneous adipose tissue during *Borrelia* replication, donated human tissues from elective surgeries were infected with 10^5^ spirochetes via subcutaneous injection and incubated *ex vivo* for 24h. DNA isolated from a portion of the tissue was used to quantify *Borrelia* loads to verify infection (Supplemental Figure S1). Skin and adipose tissue were dissected and enzymatically digested as previously described [36, 37] to generate single cell suspensions for single cell RNA sequencing. A total of 31,953 cells were isolated from the adipose stromal vascular fraction (13,023 from *Bb*-infected tissue, 18,930 from uninfected tissue) and 39,228 cells from the skin (18,356 from *Bb*-infected skin, 20,872 from uninfected skin) (Supplemental Figure S2). Clustering and Uniform Manifold Approximation and Projection (UMAP) of the sequenced cDNA libraries identified 7 major clusters in the adipose stromal vascular fraction (Figure 1A, Supplemental Figure S2A-C Bottom) and 10 major clusters in the skin (Supplemental Figure S2A-C top, S4A) defined by signature genes (Supplemental Figure S2 D-E).

**Figure 1.**
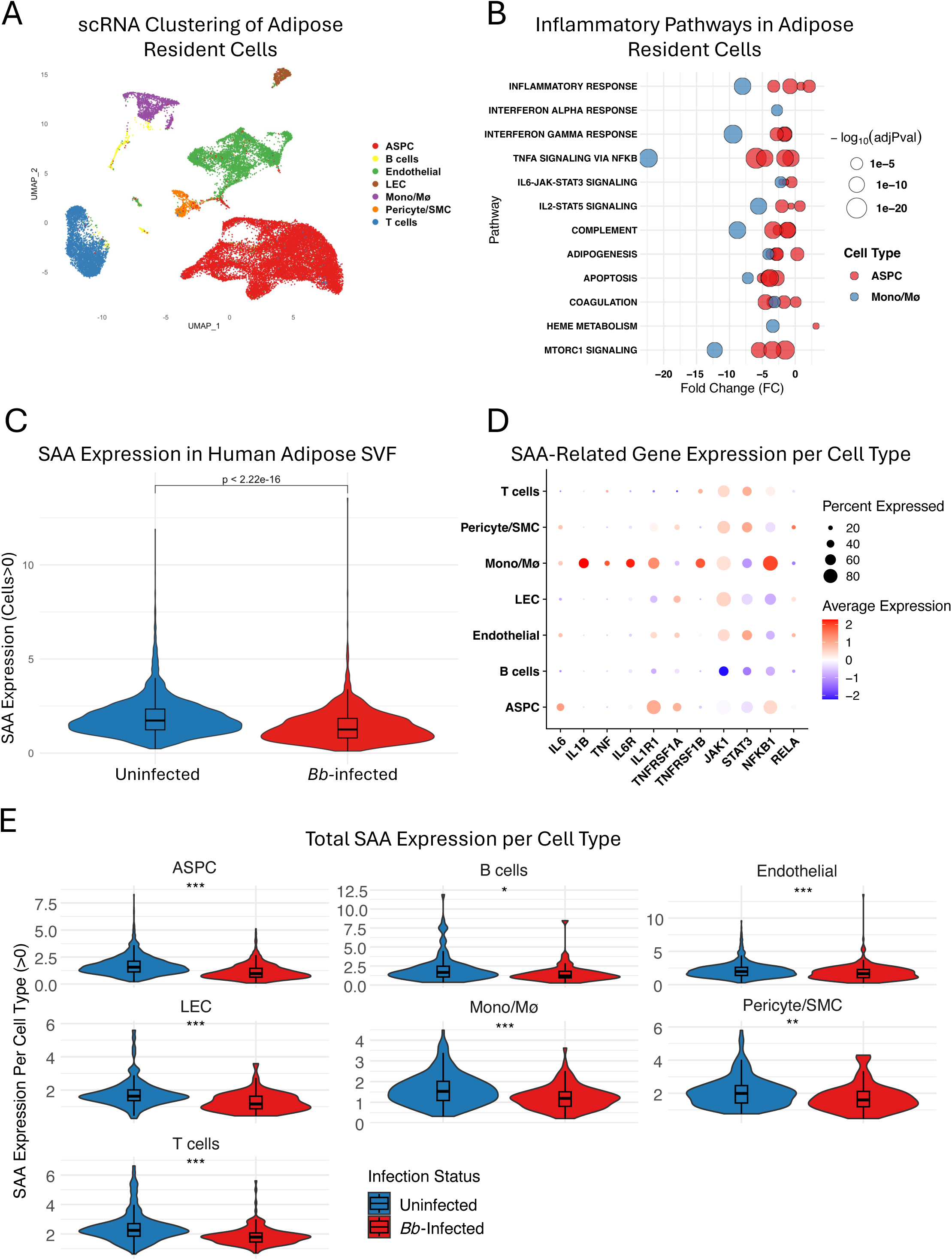
Adipose Tissue Resident Cells Show Decreased Inflammatory Expression, Including SAA Expression, After *Borrelia* Infection. Single Cell RNA Sequencing (scRNA seq) (10X Genomics) of 31,953 adipose tissue stromal vascular fraction (SVF) cells (13,023 from Bb-infected tissue, 18,930 from uninfected tissue) (n=3 Uninfected control, n=2 *Bb-*infected tissue specimens). (A) Uniform Manifold Approximation and Projection (UMAP) plots indicate 7 unique cell types with a total of 10 clusters. (B) Bubble plot shows adipose stem/progenitor cells and monocyte/macrophage (Mono/Mø) clusters and their expressed immune focused pathways from MSigDB Hallmark gene sets. (C) Total SAA (SAA1+SAA2) gene expression across tissue cohorts. (D) SAA-related genes expressed per cluster showing average expression across cell type and percent of cells expressing each gene of interest. Genes upregulated during infection were plotted in red and genes downregulated after infection plotted in blue. The size of the bubble is based on the percentage of cells within that cell type cluster expression each gene. (E) Total SAA gene expression (SAA1+SAA2) across cell clusters within each tissue cohort. Abbreviations: ASPC: adipose stem/progenitor cells; LEC: lymphatic endothelial cells; Mono/Mø: monocytes/macrophages; SMC: smooth muscle cells.

The adipose stromal vascular fraction (SVF) consists of resident immune cells, mesenchymal progenitor/stem cells, preadipocytes, and endothelial cells [38, 39]. During digestion of the tissue, fragile mature adipocytes are lysed, therefore, mature adipocytes are not included in this analysis. The adipose SVF contained a large proportion of adipose stem/progenitor cells termed (ASPCs) (precursors of mature adipocytes) [40], a cluster of monocyte/macrophages, endothelial cell populations, T cells, and pericyte/smooth muscle cells (Figure 1A). We focused on two cell types, ASPCs and adipose tissue resident myeloid cells (Mono/Macrophages), two important cell types in the adipose tissue immune response [2, 8, 41, 42]. Interestingly, we found an overall downregulation of cumulative gene expression that make up the TNF, IFN-⍺, IFN-Ɣ, and inflammatory response pathways in adipose tissue macrophages and a decrease in genes associated with adipogenesis related signaling in ASPCs (Figure 1B) in the *Bb* infected samples. ASPCs showed increased expression of the MSigDB hallmark inflammatory pathway, yet, showed downregulation of the TNFA and IL6 signaling pathways at 24h post infection in our ex vivo model system (Figure 1B). In the skin, we found elevated gene expression related to IL6 (IL6-JAK-STAT3), TNF (TNFA signaling via NFKB), IL1 (Inflammatory Response), and Type I and II interferon (IFN-⍺, IFN-Ɣ) pathways in skin resident monocytes and keratinocytes, 24h after *Borrelia* infection (Supplemental Figure S3B).

To further identify genes of biological interest within our dataset, we performed a hub gene analysis on the differentially expressed (with and without *Bb* infection) genes within our adipose tissue SVF cell clusters. We identified serum amyloid A (SAA) in ASPCs and lymphatic endothelial cells (LECs) as one of the highly ranked down-regulated genes in response to *Bb* infection within the hub gene network (Supplemental Figure S4). We also found that cumulative SAA expression across adipose SVF clusters was overall decreased after infection (Figure 1C) and was consistently downregulated across cell types including ASPCs and myeloid cells (Figure 1E). When SAA-related gene expression was examined, specifically cytokine agonists of SAA and genes expressed during SAA production [22, 23, 25, 43, 44], we observed myeloid cells had increased expression of *IL1B, TNF, IL6, IL6R, ILR1, NFKB1*, and *TNFRSF1B* (Figure 1D). Interestingly, ASPCs also showed increased RNA expression of several immune genes including *IL6, IL1R, TNF* related genes, and *NFKB1* in the setting of *Borrelia* infection (Figure 1B). Overall, we see that *Borrelia* infection modulates immune pathways in adipose tissue, including SAA which is downregulated with infection.

We also assessed skin cell clusters for genes related to SAA signaling such as cytokine agonists, agonist receptors, intermediate signaling molecules (e.g. *JAK1* and *STAT3*), and the transcription factor NFKB subunits *NFKB1* and *RELA* [22, 45]. Elevated expression levels of *IL6* were noted in skin fibroblasts, endothelial cells, and pericyte/smooth muscle cells. *IL1B* expression was also increased in innate immune cell clusters, monocytes and DC/Macrophages (Supplemental Figure S3D). We assessed total SAA expression, the combination two gene isoforms *SAA1* and *SAA2*, across skin clusters (Supplemental Figure S3C) and within each cell type specifically (Supplemental Figure S3E). Overall SAA expression was elevated in human skin after *Bb* infection predominately derived from increased expression in keratinocytes and fibroblasts.

### ASPCs Show Inflammatory and ECM-Related Interactions After B. burgdorferi Infection

To better characterize cell signaling and communication in adipose tissue after *Borrelia* infection, we utilized CellChat, a framework that infers global communication among cells using a database of ligands and receptors [46, 47]. Total predicted molecular interactions between cells show higher total cell-cell interactions in cells from infected tissue (815) compared to uninfected tissue (537) (Figure 2A) suggesting *Borrelia* infection enhances signaling within adipose tissue above homeostatic levels. To determine which cell types were involved in cell-cell interactions, we mapped the differential cell-cell interactions showing the signal strength (line thickness) from each cell type identified in adipose tissue SVF (Figure 2B). ASPCs stood out as a ‘sender’ cell with substantial autocrine and paracrine signals to LECs and monocyte/macrophage clusters suggesting an important role of ASPC signaling in subcutaneous adipose during early *Borrelia* infection.

**Figure 2.**
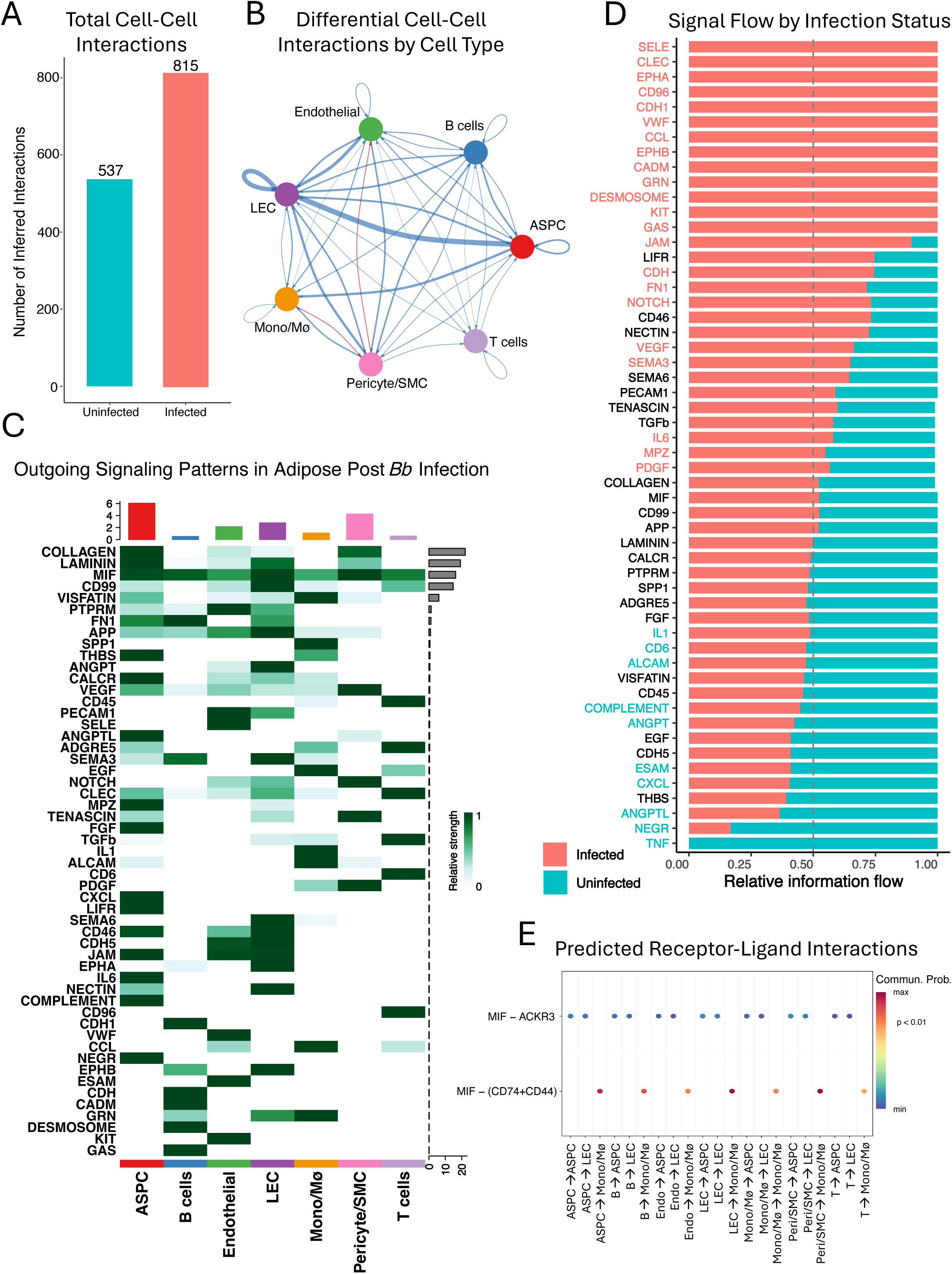
ASPCs Show Inflammatory and ECM Remodeling Interactions after *Bb* Infection. (A) Total predicted cell-cell interactions in cells from uninfected (teal color) and infected (coral color) adipose tissues using the Cell Chat ligand-receptor database. (B) A circle plot showing differential cell-cell interactions between adipose-resident cell types. Each colored node represents a specific cell type, arrows indicate ‘sender’ and ‘recipient’ for total signals, thickness of the arrow is proportional to absolute change between infected and uninfected tissues. Blue arrows indicate a decrease in signaling in the infected tissue while red represents an increase in signaling after the infection. (C) Outgoing signaling patterns computed using network centrality analysis via Cell Chat. Outgoing signal strength (top) summarizes outgoing signaling pathways (L-R systems) that are predicted to be sent out by adipose-resident cell types, corresponding with each column with cell types listed below. Each row represents a signaling pathway (i.e. collagen, laminin). The relative strength of the signal is shown via heatmap. The overall contribution of each signaling pathway across all cell types is shown with the right-side bar plot. (D) Bar plot illustrating the information flow of the top pathways across adipose tissue cell types. The total signaling strength is normalized to 1, with an information flow value of 0.5 representing no change. Genes noted in coral are upregulated during infection, and genes noted in teal are downregulated during infection. Pathways shown in colored text denote statistically significant differences according to infection status, whereas pathways in black text did not meet statistical thresholds but nevertheless exhibited modulation. (E) Predicted Receptor-Ligand (R-L) interactions specifically highlighting macrophage migration inhibitory factor (MIF) ligand visualization. Cells type indicated show an arrow from the ‘sender’ cell to the ‘receiver’ cell and the probability of communication indicated by the color with only statistically significant interactions shown. Abbreviations: ASPC: adipose stem/progenitor cells; LEC: lymphatic endothelial cells; Mono/Mø: monocytes/macrophages; SMC: smooth muscle cells.

To determine the cellular pathways that each cluster is signaling during infection, we generated the outgoing signaling patterns from cells within the infected adipose SVF (Figure 2C) and plotted the signal flow direction based on infection status (Figure 2D). Interestingly, all clusters showed a strong outgoing signal for macrophage migration inhibitory factor (MIF) (Figure 2C). ASPCs were noted as dominant broadcasters for many ECM- and immune-related signals such as MIF, IL6, complement, CD46, collagen, laminin, and fibronectin (FN1). Monocytes/macrophages were noted to send immune related signals including CCL, IL1, MIF, THBS (Thrombospondin) and ALCAM (Activated Leukocyte Cell adhesion molecule) (Figure 2C). Signal flow based on infection status (Figure 2D) shows signaling upregulated during infection (coral color) across all cell types, compared to signals downregulated during infection (teal color) normalized to a value of 1. We observed elevated IL6, MIF, collagen, fibronectin (FN1), VEGF, TGBβ, and E-selectin (SELE) signaling all increased after infection leading to pro-inflammatory, ECM remodeling, and angiogenic signals. CXCL, TNF, and IL1 signals decreased overall after infection showing a shift away from chemokine or survival signals within adipose tissue. Ligand receptor analysis of MIF showed an expected interaction with CD74/CD44 with monocytes/macrophages as the receiver cell (Figure 2E), and other cells such as ASPCs as the sender cell. Our analysis also shows an interaction with ACKR3, also known as CXCR7, and is only showing ASPCs and endothelial cells (LEC) as the receiver cells. Taken together, *Borrelia* infection amplifies cell-cell communication in adipose tissue in both total interactions and diversity of signaling pathways. ASPCs were notably found to be key hubs of proinflammatory signaling (IL6, MIF) and ECM cues (collagen, laminin, fibronectin), to surrounding immune and endothelial cells. These results suggest ASPCs act not only as structural progenitors but also as immune-active stromal cells during early *Borrelia* infection.

Despite these observed roles of adipose tissue-resident cells in our scRNAseq data, *Borrelia*-adipose interactions remain largely unexplored. To determine if adipose tissue was a site of replication for *Borrelia,* we infected C3H/HeN mice with 10^5^ *Bb* N40 spirochetes and assessed tissue loads at 7d, 14d, and 30d post infection (Supplemental Figure S5A), timepoints that correspond with early phase Lyme disease [48–51]. Flagellin gene (*FlaB*) sequences detected in tissues showed the presence of spirochete DNA in target organs of infection including skin and heart tissue, and in three different adipose depots, intrascapular, inguinal, and perigonadal adipose tissue at d14 and d30 post infection. However, live *Borrelia* cultures of individual tissue harvested at each timepoint showed inconsistent results (Supplemental Figure S5B). Cultures of injection site skin displayed live spirochetes at 7d and 14d suggesting a productive infection, but not at 30d. Mouse ears, used to detect spirochetes in a disseminated skin site, were positive at 14d and 30d post infection as expected. Adipose tissue showed live spirochetes in intrascapular adipose at 14d and 30d post infection, and a single perigonadal culture showed live spirochetes at 30d post infection suggesting dissemination into mouse adipose tissue. As we performed cardiac perfusions, removing blood vessel-resident spirochetes from tissues, these results show that *Borrelia* is present in adipose tissue depots at the timepoints studied. We can conclude that spirochetes disseminate through adipose tissue during non-hematogenous dissemination given that adipose tissue is within proximity to target organs of infection and contains many proteins which *Borrelia* spirochetes can bind, however further studies are needed to establish adipose tissue as a site of active replication.

### Serum Amyloid A2 (SAA2) binds Borrelia burgdorferi spirochetes

To better understand the function of SAA during *Borrelia* infection, we sought to determine the specific interaction between SAA and *Borrelia* spirochetes. Our single cell data indicated elevated expression in skin yet showed a downregulation in subcutaneous adipose during *Borrelia* infection. Downregulation of SAA expression would reduce SAA-*Borrelia* interactions during spirochete dissemination. Acute phase SAA has known antimicrobial properties and the ability to bind outer membrane proteins (e.g., OmpA) on Gram-negative bacteria [23–26]. SAA2 has also been previously shown to bind *Bb* N40 spirochetes incubated at 37°C using a BASEHIT library [35]. Therefore, we set out to validate these findings and determine the function of SAA binding *Borrelia* spirochetes. Whole spirochetes were incubated at 37°C for 24h to induce outer surface protein (Osp) expression consistent with replication within a mammalian host. Recombinant human His-tagged SAA2 was added to spirochetes at concentrations capable of inducing immunological activity and considered physiologically relevant corresponding to the upper limit of normal (10µg/mL) and higher acute phase reactant concentration (40µg/mL) [21, 31, 32]. Flow cytometry analysis showed a shift in fluorescence demonstrating that SAA2-His bound to N40 spirochetes as compared to control (N40 spirochetes and His-Tag Ab) or human Peptidoglycan Recognition Protein 1 (hPGLYRP1) (Figure 3A), a previously characterized protein known to bind *Borrelia* [52]. As SAA does not bind *Streptococcus pneumoniae* [32, 33], we utilized *S. pneumoniae* bacteria as a negative control in this study which also underscored the specificity of the SAA-*Bb* interaction (Figure 3B).

**Figure 3.**
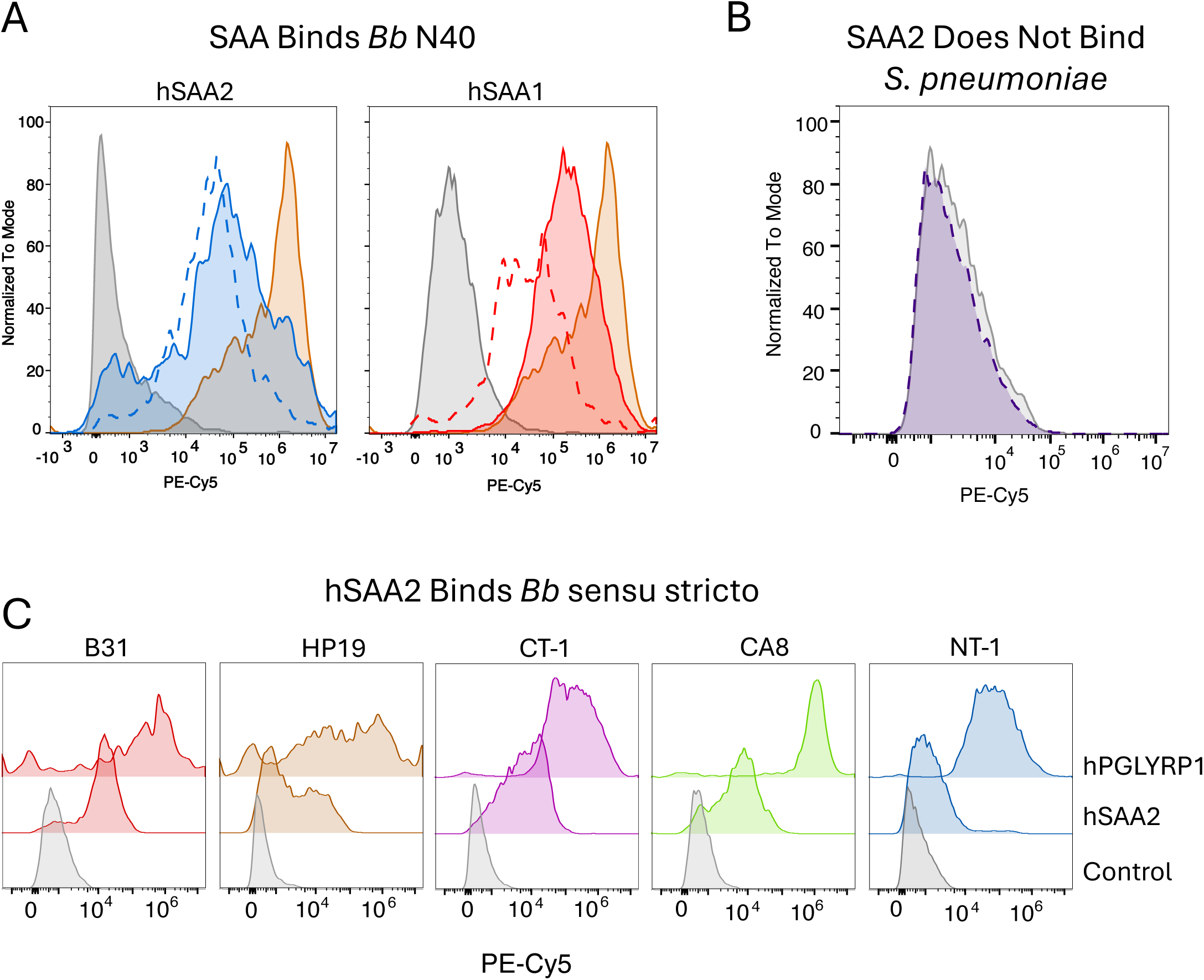
Human SAA Binds *Borrelia burgdorferi ss.* SAA2-bound spirochetes were stained for the presence of attached His-tagged protein (SAA2) detected by flow cytometry using an anti-His-tag fluorescent (PE/Cy5) conjugated antibody. (A) Histogram plots showing human SAA (hSAA2 and hSAA1), relative to non-stained controls (grey) and positive control protein (brown) known to bind *Borrelia* (Peptidoglycan Recognition Protein 1 (PGLYRP1) – Ref 39). The dashed line represents 40 µg/mL of hSAA2. The solid line represents 10µg/mL hSAA2. (B) *Streptococcus pneumoniae* incubated with 40µg/mL recombinant hSAA2 showing no increase in fluorescence compared to control. (C) Repeat binding assays showing histograms representing SAA2 binding multiple isolates of *Bb* sensu stricto (B31, CA8, HP19, CT-1, NT-1).

To determine if this binding interaction was specific to the N40 strain, we repeated experiments with relevant *Bb* sensu stricto isolates NT-1, CT-1, HP19, B31 and CA8 and showed varying binding capacity (Figure 3C). Interestingly, murine SAA2 (mSAA2) produced a larger fluorescence increase than human SAA2 (hSAA2), suggesting stronger binding of mSAA2, even at lower concentrations (Supplemental Figure S6). This supports a similar functional role for the murine ortholog in *Borrelia* infection [53]. During the acute phase response in humans, acute SAA expression is a combination of two gene products (SAA1 and SAA2) with both isotypes found the serum. With *SAA1* and *SAA2* gene isotypes sharing >95% nucleotide homology and the protein alleles sharing ∼93% amino acid sequence identity [21], it is likely that SAA1 also binds to *Borrelia* spirochetes. To determine if hSAA1 binds *Borrelia* spirochetes, repeat experiments were performed and showed hSAA1 also binds *Bb* spirochetes (Figure 3A). *Bb* N40 spirochetes bound to recombinant hSAA1 at both normal-high and acute protein concentrations. Taken together, these results validate previously identified BASEHIT library results [35] and demonstrate that both SAA1 and SAA2 bind *Borrelia burgdorferi* spirochetes.

### SAA2 Exerts Borreliacidal Effect on Bb N40 in vitro

Antimicrobial proteins, such as complement-related proteins, are a first line of defense against invading microbes. The ability of these immune mediators to bind and elicit a borreliacidal effect on invading spirochetes is paramount to generating a protective immune response. Human PGLYRP, for example, not only binds *Borrelia* but also exerts borreliacidal activity by binding newly formed *Borrelia*-derived peptidoglycan exposed during binary fission [52]. Since SAA2 is expressed at high levels, often within 24h of inflammatory stimuli [54], we sought to determine if hSAA2 can elicit a borreliacidal effect on spirochetes. *Bb* N40 spirochetes were added to clean BSKII media supplemented with increasing amounts of hSAA2 (0µg-3µg) (Figure 4). A microbial cell viability assay measuring total ATP presence showed that bacteria at 24h and 48h had decreased viability compared to cultures without hSAA2 protein. These results indicate hSAA2 binding to *Borrelia* spirochetes leads to decreased viability at timepoints consistent with acute phase SAA expression in humans.

**Figure 4.**
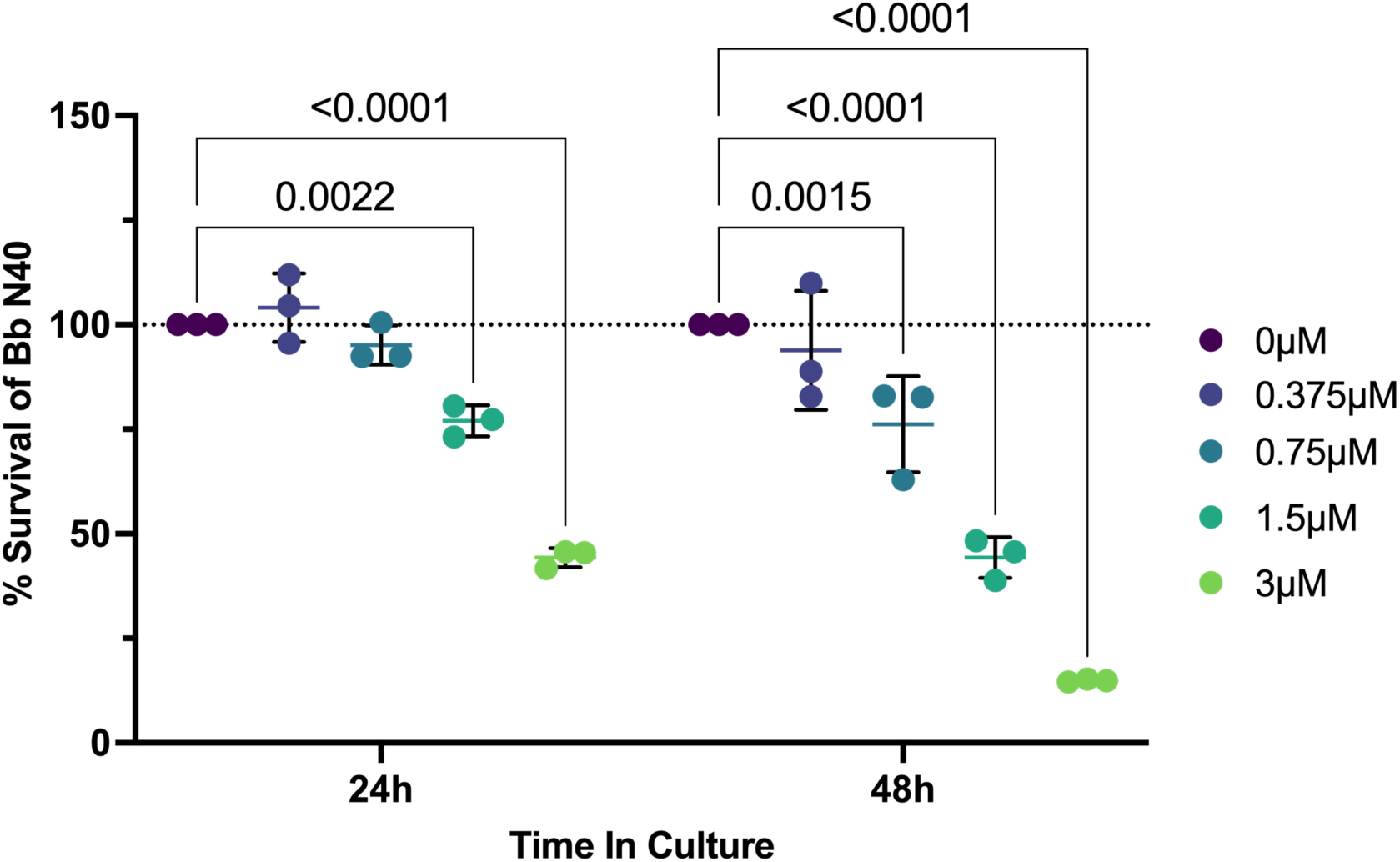
Human SAA2 has Borreliacidal Effect in a Dose-Dependent Manner. *Bb* N40 spirochetes were added to clean BSKII media supplemented with no protein (control) or increasing concentrations (0µM-3µM) of recombinant hSAA2. Cultures were sampled at 24h and 48h post incubation. Bacterial abundance was assessed using the BacTiter Glo reagent measuring luminescence of total bacterial-derived ATP in the culture. Values plotted are luminescence values normalized as survival percentages of *Bb* + 0µM hSAA2 (1x PBS) in culture. Statistical significance was determined using a two-way ANOVA followed by post hoc Dunnett’s test comparing treatment groups to the control (0µM). Only significant p values are shown. Data is combined of two independent experiments performed in triplicate.

### SAA2 Enhances Phagocytosis of Bb Spirochetes in-vitro

Prior studies demonstrate SAA’s ability to bind and opsonize bacterial pathogens, facilitating phagocytosis of gram-negative bacteria such as *E. coli, P. aeruginosa, V. cholerae, K. pneumoniae,* and the acid-fast bacterium *M. tuberculosis* [31–33]. Because serum SAA concentrations peak ∼24h after injury or infection [54], these data highlight a key role for this acute phase protein in binding/opsonizing bacteria to promote phagocytosis before protective antibodies develop during a primary infection. To determine if SAA2 opsonizes *Borrelia* to enhance phagocytosis, CFSE-stained *Bb* N40 spirochetes preincubated with SAA2 were added to membrane-labeled (DiD) PMA-differentiated THP-1-macrophages at a multiplicity of infection (MOI) equal to 10 (Supplemental Figure S7A). Flow cytometry was used to measure double positive populations which represent co-localization of phagocytic cells and SAA2-bound spirochetes relative to stained control cells or spirochetes (Supplemental Figure S7B). We observed an increase of double positive cells over time and in a dose-dependent manner (Figure 5A, S7C) and colocalization of spirochetes with THP-1 cell membranes (Figure 5B). With 1µg/mL of recombinant hSAA2, membrane-stained cells that also stained positive for *Bb* increased from 1.27% at 0.5h to 4.98% at 1 hour, 12.1% at 2 hours, and 30.5% at 3 hours of co-culturing. Increasing hSAA2 concentration to 10µg/mL resulted in 65.6%, 62.9%, 74.2%, and 80.7% double positive cells at 0.5h, 1h, 2h, and 3h, respectively (Figure 5A, Supplemental Figure S7C). Normal human serum (NHS) was employed as a control, since it contains complement components that mediate opsonization of bacterial pathogens [55]. Repeat experiments using recombinant mSAA2 showed a steady increase of the double positive population over time, representing phagocytosis, however, the percentages did not reach statistical significance (Supplemental Figure S8). It is possible that mSAA2 is less effective at opsonization with human phagocytes in vitro. Taken together, these results indicate that hSAA2 binding *Borrelia* spirochetes results in opsonization and facilitates phagocytosis in vitro.

**Figure 5.**
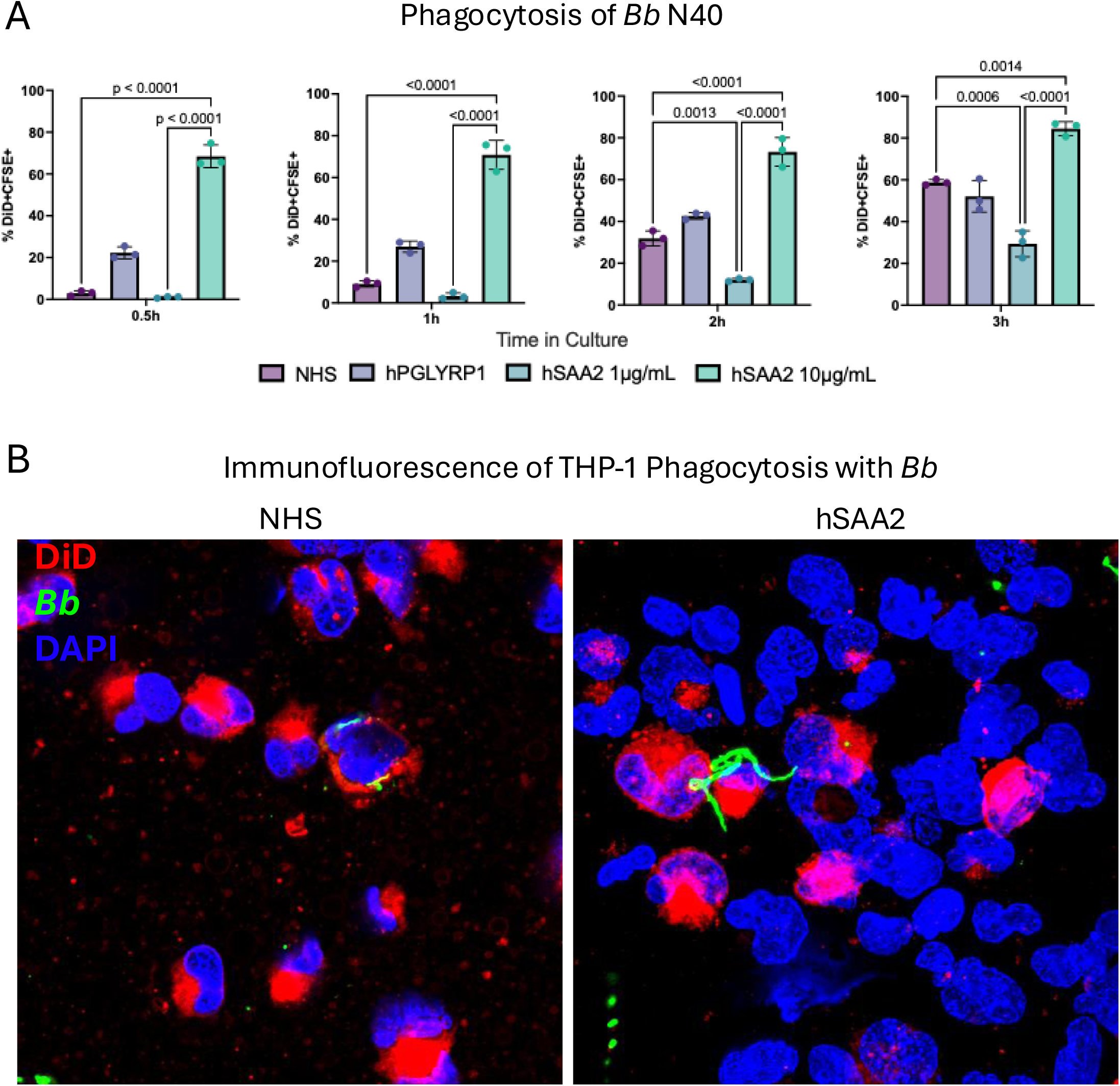
SAA2 Opsonizes *Bb* Spirochetes and Enhances Phagocytosis. (A) DiD stained THP-1 differentiated macrophages were incubated with CFSE-labeled N40 spirochetes for 0.5h, 1h, 2h, or 3h at a multiplicity of infection equal to 10 (MOI:10), and hSAA2 at 1 ug/mL and 10 ug/mL. Graphs represent percentages of DiD+CFSE+ double positive populations of cells quantified by flow cytometry. Statistical significance was determined a two-way ANOVA followed by post hoc Tukey test to correct for multiple comparisons. (B) Immunofluorescence of DiD-labeled differentiated THP-1 macrophages and CFSE-labeled *Bb* at 2h post co-culture at 10µg/mL. Abbreviations: NHS: Normal human serum. PGLYRP1: peptidoglycan recognition protein 1, DiD(red)= cell membrane, *Bb* (green) = *Borrelia Burgdorferi*, DAPI (blue) = nuclei.

## Discussion

We used single cell RNA sequencing to evaluate the adipose tissue immune landscape during *Borrelia* infection. Overall, we found that *Borrelia* modulates immune pathways in adipose tissue, in addition to skin. During infection, we observed downregulation of immune pathways (Inflammatory Response, IFN-α and IFN-γ responses, TNF signaling, IL2-STAT5 signaling, Complement) and downregulation of SAA in monocytes/macrophages (Figure 1). SAA1 also emerged as an important hub gene in both ASPCs and LECs (Supplemental Figure S4). As expected, we found upregulation of most immune pathways (Inflammatory response, IFN-α and IFN-γ responses, TNF signaling, Complement) in both keratinocytes and monocytes in the skin (Supplemental Figure S3). SAA was notably upregulated in skin cells, mostly in keratinocytes and fibroblasts. ASPCs from infected adipose tissue emerged as signaling hubs expressing outgoing signals for ECM-related (collagen, laminin, FN1) and immune-related (MIF, CXCL) signaling pathways highlighting an immune-active role of ASPCs during *Borrelia* infection (Figure 2). Specific inflammatory genes such as IL1B, TNF, IL6R, and NFKB1 were upregulated with infection in adipose tissue monocytes/macrophages. ASPCs were the only cell type that showed overall upregulation of the inflammatory response pathway, with specific upregulation of IL6, IL1R1 and NFKB1 (Figure 1).

Lyme Disease patients mount an acute phase response to early *Borrelia burgdorferi* (*Bb*) infection with increased levels of acute phase response proteins including serum amyloid A (SAA) [19]. Previous studies have shown that, SAA2 can bind *Bb* N40 spirochetes using BASEHIT screenings [35]. We further validated the binding interaction between human SAA and *Borrelia* spirochetes to determine the functional implications of this host-pathogen connection. Specifically, we found that human SAA can bind *Borrelia* spirochetes (Figure 3), validating prior BASEHIT library results, we demonstrated that hSAA2 bound to whole spirochetes exerts a borreliacidal effect on bacterial cultures at 24h and 48h (Figure 4) and finally, SAA2 enhances phagocytosis of *Borrelia* spirochetes *in vitro* (Figure 5).

SAA is known to be an inducer of the inflammasome and contributes to downstream differentiation of the Th17 response [56]. Within adipose tissue SAA is a pro-inflammatory adipokine that induces the production of pro-inflammatory cytokines [57]. Additionally, SAA expression inhibits adipogenesis and contributes to insulin resistance [58]. Within our own data we saw downregulation of the adipogenesis hallmark pathway in ASPCs (Figure 1B). Therefore, we hypothesize that *Borrelia* is downregulating SAA for its benefit. Lactobacillus downregulates SAA in gut epithelium which in turn impairs Th17 differentiation to exert a protective effect in the setting of colitis [59]. A recent study showed that epiploic white adipose tissue (adipose tissue adjacent to colon) harbors high proportions of SAA expressing adipocytes along with several leukocyte populations. This work demonstrates that expression of SAA in adipocytes is induced by inflammatory signals such as lipopolysaccharide (LPS), and that SAA then activates downstream immune responses in adipose tissue resident cells [60]. Downregulation of immune pathways, including SAA by *Borrelia* could create a sanctuary in adipose tissue and allow persistence of this pathogen in adipose tissue or very simply allow it to further disseminate throughout the body (i.e., through inhibition of phagocytosis). Further investigation is needed to understand how *Borrelia* is inducing the downregulation of SAA in adipose tissue.

Skin is a natural replication site for spirochetes during infection and is directly adjacent to subcutaneous adipose tissue, a complex multicellular organ with immune, metabolic, and endocrine function [2, 8, 61] that is involved in the pathogenesis of several vector borne pathogen infections [10, 12, 15, 16]. During *Borrelia* infection in the mammalian host, a myriad of surface proteins expressed on the spirochete surface [62] play critical roles in allowing *Borrelia* to actively migrate from the site of infection in the skin to distal organs, collagenous tissues, joints, and synovial fluid of the host [6]. By investigating immune signaling in adipose tissue 24h post *Borrelia* infection, we identified several potential interactions in our network analysis that are worth future exploration. We observed a strong outgoing signal for laminin in ASPCs, LEC, and pericyte/SMC in our cell chat analysis. Complexes of SAA and *Borrelia* could have the ability to bind laminin. This is consistent with SAA’s capacity to bind both ECM components, and microbial surfaces [31, 33]. Several *Borrelia* outer surface proteins, including DbpA/B and BB0406 directly bind laminin to promote tissue colonization and dissemination. SAA is also known to bind laminin [63, 64] which has been shown to facilitates mast cell and T cell adhesion [65, 66], and thus provides a bridging point for meaningful interactions. Such bridging interactions may not only restrict *Borrelia* dissemination but also amplify local inflammation through activation of SAA-responsive receptors (TLR2/4, FPR2) on stromal and immune cells. A strong outgoing signal for MIF was also noted in all cell types (Figure 2C) during infection. To date, the role of MIF during *Borrelia* infection has not been described in detail. As the name suggests, MIF regulates effector immune cell migration to promote their persistence at sites of infection and expression of inflammatory cytokines [67]. West Nile virus (WNV) patients showed elevated MIF levels compared to controls with levels even higher in patients that succumb to disease [68]. MIF knockout mice showed decreased lethality of neuroinvasive disease compared to WT controls during WNV infection[68]. Ligand receptor analysis of MIF shows an expected interaction with CD74/CD44 with monocytes/macrophages as the receiver cell (Figure 2E), and an interaction with ACKR3 (CXCR7) with ASPCs and endothelial cells as the receiver cells. Other studies have shown the MIF-CD74/CD44 interaction to play a role in wound healing and the wound microenvironment [69, 70], and in the context of cancer, the MIF-ACKR3 interaction has recently been shown to drive repression of adipogenesis in ASPCs [71], a potential interaction mechanism behind the observed reduction in adipogenesis pathway signaling reported here (Figure 2D, E). Continued investigation into the role of MIF and *Borrelia* pathogenesis is needed to determine its role during spirochete dissemination through adipose tissue.

Non-hematogenous dissemination of *Borrelia* (direct migration through tissues or motility driven pathogenesis) is an important aspect of *Borrelia* tissue invasion. Seven to eleven periplasmic flagella enable the spirochetes to actively migrate ∼4µm per second [72] along the plane of the epidermis and medially towards the dermis microvasculature and subcutaneous tissue causing erythema migrans [3]. *Borrelia* spirochetes are known to bind integrins ⍺5β1 and ⍺Vβ3 [73, 74], both expressed by adipocytes and ASPCs [75, 76] which make *Borrelia* dissemination through adipose tissue a likelihood. However, there are limited studies that have evaluated *Borrelia* infection within adipose tissue. Histopathologic analysis of perisynovial adipose tissue from rhesus macaque stifle joints in a late disseminated *Borrelia* infection demonstrated positive fluorescent staining for *Borrelia* during late-stage arthritic disease [9]. Infection of this adipose tissue depot may coincide with the arthritic symptomology seen during the late disseminated phase of Lyme disease. Although this dissemination into perisynovial adipose has not been reported in humans, panniculitis, inflammation of the subcutaneous fat, has been reported as a manifestation of Lyme borreliosis [77–80]. Finally, studies that evaluated diet-induced obese mice in the setting of *Borrelia* infection found that obese mice displayed more severe Lyme carditis and delayed polymorphonuclear cell infiltration [81],and had attenuated *B. burgdorferi*-specific IgG production within 8 weeks of infection [82] demonstrating adverse shifts in immunity when obesity is present in the host. Therefore, adipose tissue in the context of *Borrelia* infection is an unexplored area of research that deserved further investigation.

Our study is the first to really evaluate *Borrelia* infection in the adipose tissue microenvironment. We were able to demonstrate with our data that *Borrelia* was detected in human skin and adipose tissue explants 24 hours post infection (Supplemental Figure S1). Furthermore, *Borrelia* infection of C3H/HeN mice demonstrated that *Borrelia* was detected in adipose tissue depots on d14 and d30 (Supplemental Figure S5A) timepoints that correspond with early phase Lyme disease [48–51]. Live *Borrelia* cultures of adipose tissue showed live spirochetes in intrascapular adipose at 14d and 30d post infection, and a single perigonadal culture showed live spirochetes at 30d post infection suggesting dissemination into mouse adipose tissue. Overall, these results show that *Borrelia* is present in adipose tissue depots. Adipose tissue is a known reservoir for vector-borne parasites like *Trypanosoma cruzi, T. brucei*, and *Plasmodium* [10–12]. Further investigation is needed regarding the role *Borrelia* is playing in adipose tissue.

Several questions remain. The specific mechanism by which SAA binds *Borrelia* remains to be determined. Many of the *Borrelia* lipoproteins contain lipid moieties that anchor the protein into the outer membrane of the spirochete. It is possible the hydrophobic anchor of *Borrelia* lipoproteins associates with SAA hydrophobic alpha helices similar to the SAA-HDL interaction [34, 54]. Alternatively, electrostatic interactions may facilitate SAA binding to acidic residues on *Borrelia* lipoproteins with SAA glycosaminoglycan-binding domains, as is the case for acidic loops on OspC or DbpA/B, or GAG-like regions on Bgp [83, 84], therefore mimicking the SAA-GAG interaction. Further investigation is required to determine which, or if both mechanisms are used. Finally, we found that SAA exerted a borreliacidal effect on bacterial cultures. Increasing concentrations of recombinant SAA ranging from 1.5-3.0µM at 24h and from 0.75µM to 3µM at 48h in culture in culture showed a negative effect compared to control treated *Borrelia* (Figure 4). The mechanism by which SAA exerts this decline in spirochete viability is unclear. However, it has been reported that an SAA isotype (SAA1.1) can form voltage-dependent ion channels in lipid bilayers [85] suggesting a potential mechanism for SAA pathogenesis.

Overall, we report here that *Borrelia* infection leads to the modulation of several immune pathways in adipose tissue, including the downregulation of SAA. We found SAA binds, enhances phagocytosis, and exerts a borreliacidal effect on *Borrelia* spirochetes. These findings highlight an underappreciated role for adipose tissue in the local host response to *Borrelia* spirochetes and underscores the need for continued characterization of this complex organ in the context of Lyme disease.

## Methods

### Ethics Statement

C3H/HeN mice were obtained from Charles River Laboratories. All mice were housed and maintained at the Yale University School of Medicine Animal Resource Center under approved protocols (Protocol 07941) according to Yale University Institutional Animal Care and Use Committee (IACUC) and in accordance with the American Association for Accreditation of Laboratory Animal Care (AAALAC) guidelines. Human subjects undergoing elective surgeries, such as breast reconstruction surgery, donated tissues used for the human explant model through a protocol approved by the Human Research Protection program of the Yale School of Medicine. Informed consent was obtained from all subjects prior to participation. Exclusion criteria included active cancer and treatment, any active infection, and use of anticoagulation.

### Bacterial and Mammalian Cell Culture

*Borrelia burgdorferi* spirochetes (N40, B31, HP19, CA8, NT-1, CT-1), provided by E.F., were cultured in Barbour Stoenner-Kelly (BSK) II complete media supplemented with 6% rabbit serum at 33°C. Live spirochete density was determined by dark field microscopy (40x) using a Neubauer improved hemocytometer. Spirochetes were incubated at 37°C for 24h minimum prior to assays with SAA to ensure mammalian specific Osp expression on the spirochete surface. *Streptococcus pneumonae* Serotype 4 (NCBI BioSample: ABC020012001) was kindly provided by Dr. Inci Yildirim (Yale) and cultured in Brain Heart Infusion broth (Millipore Sigma) at 37°C + 5% CO_2_. Optical density was quantified using a Thermo Scientific BioMate 160 cuvette spectrophotometer. Mammalian THP-1 cells were cultured in complete RPMI 1640 media supplemented with 10% fetal bovine serum and 1% penicillin-streptomycin and incubated at 37°C + 5% CO_2_. Cell density was determined using a Bio-Rad TC-20 automated cell counter with trypan blue to determine cell viability.

### Binding Assay, Borreliacidal Assay, and Phagocytosis Assays

A total of 2×10^7^ spirochetes, or equivalent number of streptococci, were used for each binding assay in triplicate. Spirochetes were pelleted at 5000 x *g* for 5 minutes, washed with 1x PBS, then resuspended in solution with SAA2 recombinant protein (hSAA2-His: MedChemExpress HY-P70985, Elabscience PKSH033532; mSAA2-His: antibodies online ABIN7088660) at 1µg/mL, 10µg/mL or 40µg/mL for 1h at room temperature. After recombinant protein was allowed to bind, spirochetes were washed and resuspended in 1x PBS. His-tagged recombinant proteins bound to spirochetes were labeled using a PE-Cy5 conjugated anti-His-tag antibody (Biolegend 362635) for detection by flow cytometry [52] using a Beckman Coulter Cytoflex LX. Human PGLYRP1 (R&D Systems 2590-PGB) [52] was used as a positive control and allowed to bind to spirochetes as stated above at 40µg/mL.

To determine the effect of SAA2 on live spirochetes, 5×10^6^ spirochetes were resuspended in fresh BSKII media adjusted to contain increasing concentrations of SAA2 at 0µM (control), 0.375µM, 0.75µM, 1.5µM, and 3.0µM in solution. Spirochetes were incubated at 37°C with sample aliquots harvested at 24h and 48h timepoints. 20µL aliquots taken from each replicate at each timepoint were mixed with the BacTiter-Glo reagent (Promega G8230) 1:1 and measured luminescence on a Tecan infinite M200 Pro plate reader using a black-clear bottom 384-well plate format.

A total of 5×10^5^ THP-1 monocyte-like cells were seeded into a 24-well plate and stimulated with 1µM phorbol myristate acetate (PMA) overnight to induce an adherent macrophage-like cell phenotype. Differentiated cells were stained with Vybrant DiD cell-labeling solution (Invitrogen V22887) according to the manufacture’s specifications. SAA2-bound spirochetes were then stained with carboxyfluorescein succinimidyl ester (CFSE) (eBioscience 65-0850). Spirochetes were added to THP-1 cells at a multiplicity of infection (MOI) equal to 10. Flow cytometry was used to determine the total number of DiD-positive CFSE-positive cells.

### Human Tissue Explant Infections and Single Cell RNA Sequencing

Tissue specimens (n=3 Uninfected control, n=2 *Bb-*infected tissue specimens) were cut into ∼1-2cm^2^ cubes that contained epidermis, dermis, and subcutaneous adipose tissue [86]. Each sample was incubated in Netwell inserts with 115mm diameter and a 74µm mesh pore size (Corning). Samples were inoculated subcutaneously with either 10^5^ live spirochetes or clean BSKII media adjusted to a 50µL total injection volume. Netwells were incubated in 12-well plates with 1mL RPMI 1640 supplemented with 10% FBS without antibiotics at 33°C + 5% CO_2_. After 24h incubation, skin and adipose were separated and enzymatically digested as previously described [36, 37]. Skin was incubated in RPMI 1640 medium containing 5% FBS and dispase II (10mg/mL) for 45min at 37°C in a shaking water back incubator set at 200rpm. The skin was removed, and mined using a scalpel, and added to RMPI 1640 containing 5% FBS, 0.5mg/mL Liberase TM (Millipore Sigma), and 30 U/ml DNase I (Millipore Sigma) and incubated for 45min at the previous conditions. Resulting cell suspensions were passed through a 100µm, 70µm and 40µm filters to remove debris, washed, and resuspended in 1x PBS + 20% FBS to maintain cell viability. Cell suspensions volumes were adjusted to be added to the Chromium Next GEM Single Cell 3’ kit (10x Genomics) at maximal concentrations.

Adipose tissue was processed by mincing with a scalpel and adding the tissue to a dissociation solution made of Krebs Ringer Buffer (KRB) containing 0.8mM zinc chloride (ZnCl_2_), 1mM magnesium chloride (MgCl_2_), and 1.2mM calcium chloride (CaCl_2_) with collagenase II dissolved at 1mg/mL and 5% BSA. Tissue suspension was incubated in a shaking water bath (200 RPM) at 37°C for 30-45min until suspension was homogenous. Suspension was passed through a 100µm and 70µm sieves, with 1 volume of KRB was passed though the sieves to ensure maximum cell recovery. Cell suspension was centrifuged at 600 x *g* for 6 min to pellet the stromal vascular fraction (SVF) cells. The yellow oily floating lipid layer from lysed adipocytes was aspirated off and stored at −80°C. The remaining supernatant was discarded. Cells were resuspended in fresh complete RPMI 1640 media and red blood cells were removed using 1x RBS lysis buffer (eBioscience). Remaining SVF cells were pelleted, resuspended in complete RPMI, and counted on a hemocytometer using trypan blue. SVF cells were added to the Chromium Next GEM Single Cell 3’ kit (10x Genomics) at maximal concentrations.

### Single cell RNA sequencing analysis

cDNA libraries were generated from either skin or adipose SVF using the Chromium Next GEM Single Cell 3’ kit reagents, Illumina workflow, on a NovaSeq platform. The 10x Cell Ranger software was used to align reads, process unique molecular identifiers (UMIs), and generate the expression count matrix. We utilized a quality control pipeline as described in the Seurat algorithm, using only genes detected in at least 3 cells and cells with at least 200 genes detected in the analysis [87]. We filtered out cells with mitochondrial DNA content higher than 10% and excluded predicted doublets using the default Scrublet software package settings [88]. The resulting expression matrix was normalized using Seurat (v4.3.0) and corrected for batch effect using Seurat’s canonical correlation analysis (CCA) method [87]. Clustering and integration was performed using Louvain algorithm after Seurat’s SCTransform [89]. Cell type annotation was performed using Seurat’s ‘FindConvervedMarkers’ function and comparing generated marker tables to identifying cell markers from single cell tissue atlases [37, 90–94]. To determine expression contrasts between infected and uninfected cohorts, differential gene expression analysis was performed using the model-based analysis of single-cell transcriptomics (MAST) algorithm [95], using a log fold-change of 0.5 with an adjusted P value of 0.05 to find significant differential expressed genes. Pathway analysis per cluster was performed using the single cell pathway analysis (SCPA) tool [96] using MSigDB Hallmark pathways, Kyoto Encyclopedia of Genes and Genomes (KEGG) pathways, and REACTOME pathways. To determine cell-cell interactions analyses, CellChat (v1.6.1) was used to determine possible receptor-ligand (R-L) interactions using the built in tools for visualization [46, 47]. All analysis was performed using R Studio (2025.05.1+513), Jupyter Notebook (v4.4.6), and Python (v3.13).

### Borrelia infection in C3H/HeN mice

Four- to six-week-old female mice were anesthetized using isoflurane and infected with 10^5^ spirochetes (50µL total volume) using an intradermal inoculation along the dorsal thoracic region of the mouse (scruff) using a 30ga insulin syringe. Uninfected control mice were inoculated with clean BSKII media. Mice were euthanized at 7d, 14d, 30d post infection using carbon dioxide according to approved protocols. A cardiac perfusion was performed with 20mL of 1x PBS containing 20 U/mL heparin injected into the left ventricle to remove the possibility of spirochetes residing in blood vessels to give a false positive signal in qPCR of tissue cultures. Tissues harvested included skin, heart, and a large joint, along with intrascapular, inguinal, and perigonadal adipose depots [97, 98].

### DNA isolation and FlaB quantification from human and mouse tissues

Harvested tissues, from human explants or mouse tissues, were homogenized and incubated in tissue lysis buffer overnight at 56°C according to the Qiagen DNeasy Blood and Tissue kit according to the manufacturer’s specifications. A Nanodrop 2000 (Thermo Scientific) was used to check DNA concentration and purity. Absolute quantification of *FlaB* sequences (forward primer: 5’-TTCAATCAGGTAACGGCACA-3’, reverse primer: 5’- GACGCRRGAGACCCTGAAAG-3’) within tissues was performed using 50ng of sample DNA and the iTaq Universal SYBR Green Supermix (Biorad) on a CFX96 Touch quantitative thermocycler (Biorad). Cycle threshold values from samples were compared against a standard curve generated by serial dilutions of DNA from a stock of *Borrelia* spirochetes with a known concentration, normalized by total mass of template nucleic acid in the reaction.

## Acknowledgements

We would like to acknowledge and thank all our human subjects that donated their surgical tissue for this study. The sequencing for this study was conducted by Dr. Mei Zhong at the Yale Stem Cell Center Genomics Core facility. We would also like to acknowledge Dr. Inci Yildirim for donating the *Streptococcus* bacteria used in this study. We are also grateful to Dr. Albert Shaw and Dr. Sukanya Narasimhan for their thoughtful feedback on this manuscript. Finally, we would like to thank members of our HHMI EPI consortium for all their input and feedback, especially Dr. Photini Sinnis and Dr. Maudry Laurent-Rolle.

## Funding

This work was funded by the HHMI Emerging Pathogens Initiative.

---

**Supplemental Figure S1.**
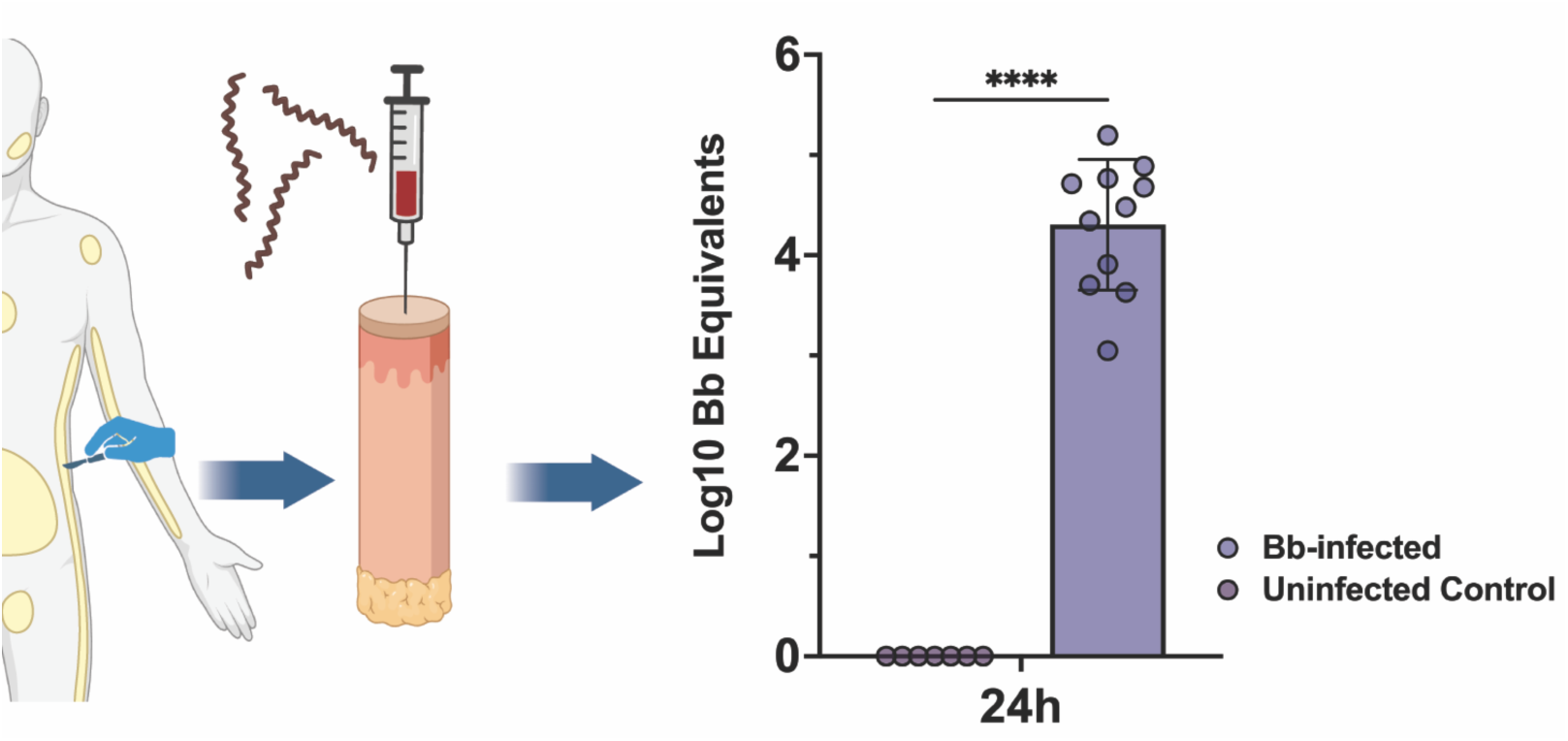
FlaB Detection in Human Tissue DNA. Donated human tissue was cut into 1cm^3^ samples and subcutaneously inoculated with 10^5^ spirochetes. At 24h post infection, a small portion of the human tissue was removed, homogenized, and used for total DNA isolation. Quantification was performed using a standard curve generated from serial dilution of DNA isolated a known number of spirochetes. Cycle threshold (Cq) values plotted as a function of Log10 spirochetes was used to determine the bacterial loads in tissue from infected samples relative. *Bb* infected n= 11, Uninfected control n=7. Statistical significance was performed using a paired T test. **** indicates a P values <0.0001.

**Supplementary Figure S2:**
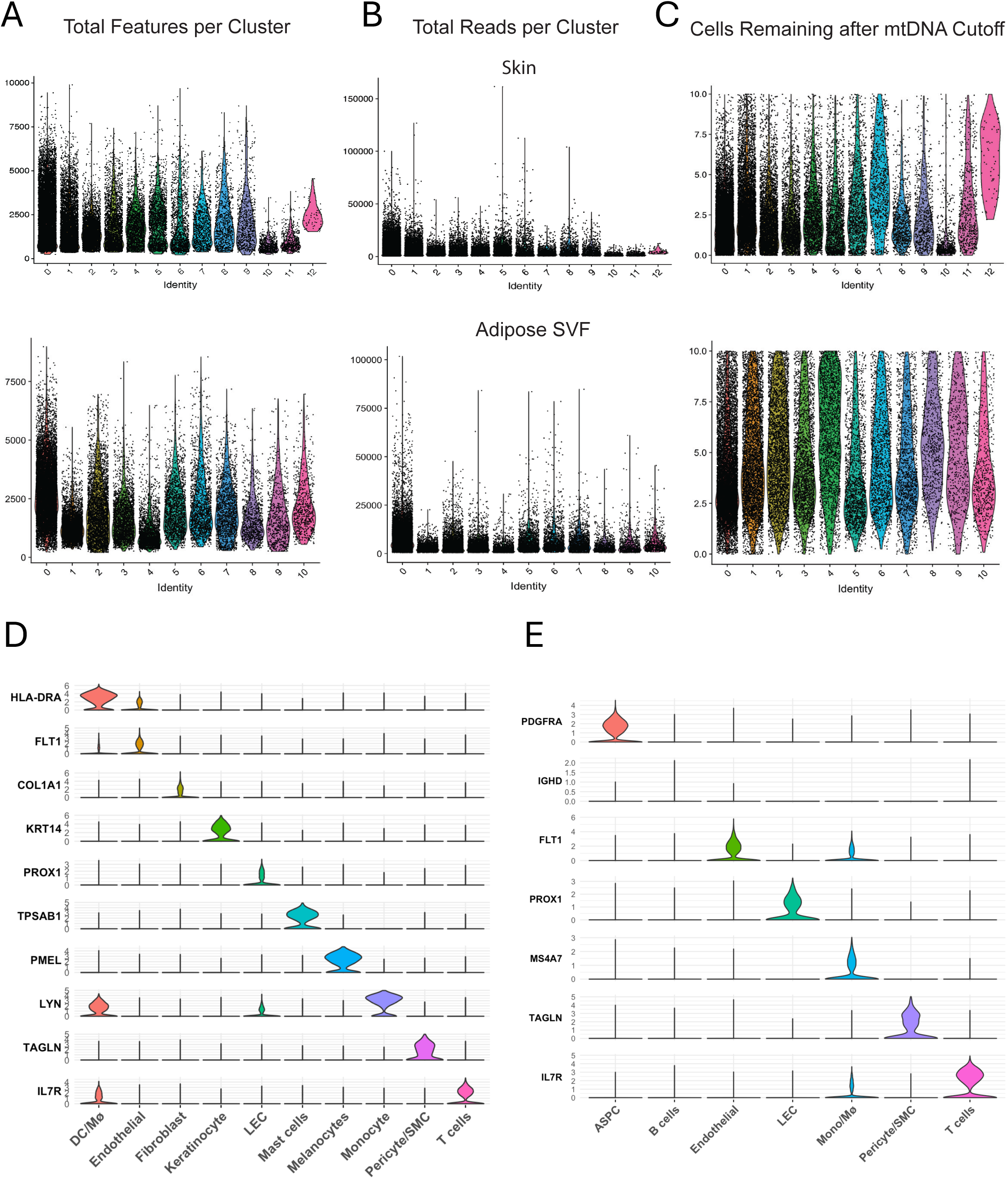
Total Features, Reads, and Identified Marker Genes from Human Skin and Adipose Cells after Single Cell RNA Sequencing. (A) Total features for human skin (top row) and adipose (bottom row) samples, (B) reads per cluster for human skin (top row) and adipose (bottom row) samples, and (C) cells remaining after single cell quality control and mitochondrial DNA cutoffs (10%) for human skin (top row) and adipose (bottom row) samples. (D-E) Violin plots of individual marker genes used to identify skin (D) resident populations and (E) adipose resident populations. Abbreviations: ASPC: Adipose stem/progenitor cells, LEC lymphatic endothelial cells, Mono/Mø: monocytes/macrophages, SMC: smooth muscle cells. (A-C) Adipose clusters: Cluster 0: ASPC(1); Cluster 1: T cells; Cluster 2: Endothelial(1); Cluster 3: ASPC(2); Cluster 4: Endothelial(2); Cluster 5: ASPC(3); Cluster 6: ASPC(4); Cluster 7: Monocyte/Macrophage; Cluster 8: SMC/Pericyte; Cluster 9: B Cell; Cluster 10: LECs. Skin clusters: Cluster 0: Fibroblast(1); Cluster 1: Endothelial(1); Cluster 2: T cells; Cluster 3: LECs; Cluster 4: Fibroblast(2); Cluster 5: Dendritic cells/macrophages; Cluster 6: Keratinocytes; Cluster 7: Melanocytes; Cluster 8: Endothelial(2); Cluster 9: Pericyte/SMC; Cluster 10: Monocytes; Cluster 11: Mast Cells; Cluster 12: Endothelial(3). (D-E) Multiple ASPCs, fibroblasts, or endothelial cell clusters were combined within each tissue and represented by a single cell type (i.e. ASPCs in adipose tissue is represented by ASPC clusters 0, 3, 5, and 6 in adipose tissue). Please note that the signature gene for B cells in Adipose tissue was IGHD, yet this cluster has significantly low cell counts compared to other clusters.

**Supplementary Figure S3:**
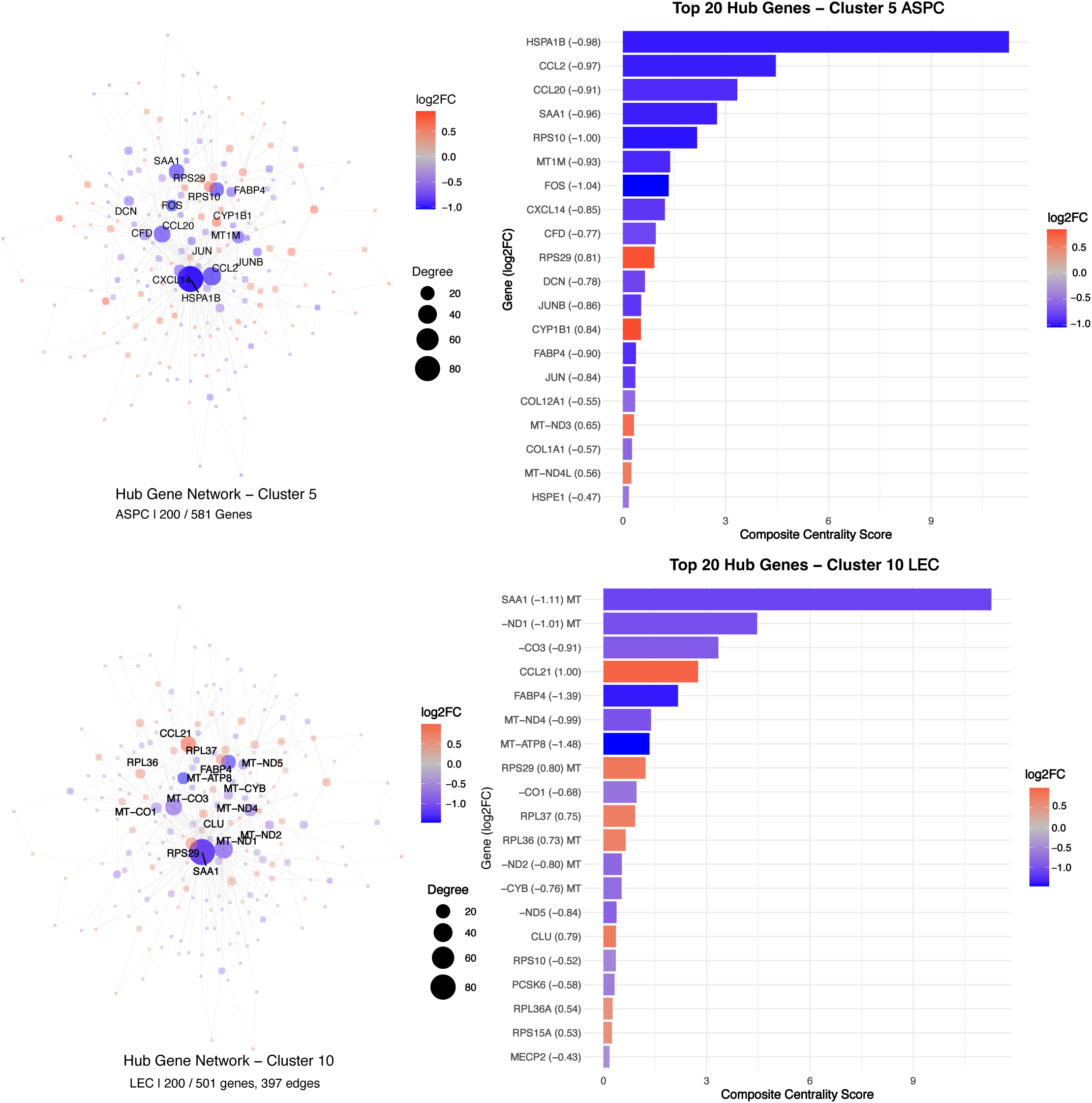
Single Cell Sequencing Analysis of *Bb*-Infected Human Skin. (A) Clustering using Uniform Manifold Approximation and Projection (UMAP) showing 10 unique cell types from 12 clusters from a total of 39,228 cells from 2 infected and 2 uninfected skin specimens with adipose removed (18,356 from Bb-infected skin, 20,872 from uninfected skin). (B) Inflammatory-specific pathways modulated in skin-resident DC/macrophages, keratinocytes, and monocytes. Normalized expression data for each cluster was used as a query to determine differentially expressed MSigDB Hallmark pathways. (C) Total SAA expression in cell from both control and *Bb-*infected human skin. Only cells with expression above zero are shown. (D) Average expression of genes related to SAA expression across skin cell types. Expression is normalized to uninfected control genes showing upregulation in infected tissue was red and downregulation after infection in blue. Size of the bubble corresponds to the percentage of each cluster expressing each gene. (E) Violin plots showing total SAA expression per cell type within uninfected control skin cells (blue) and *Bb*-infected skin cells (red). Abbreviations: ASPC: adipose stem/progenitor cells; LEC: lymphatic endothelial cells; Mono/Mø: monocytes/macrophages; SMC: smooth muscle cells.

**Supplemental Figure S4.**
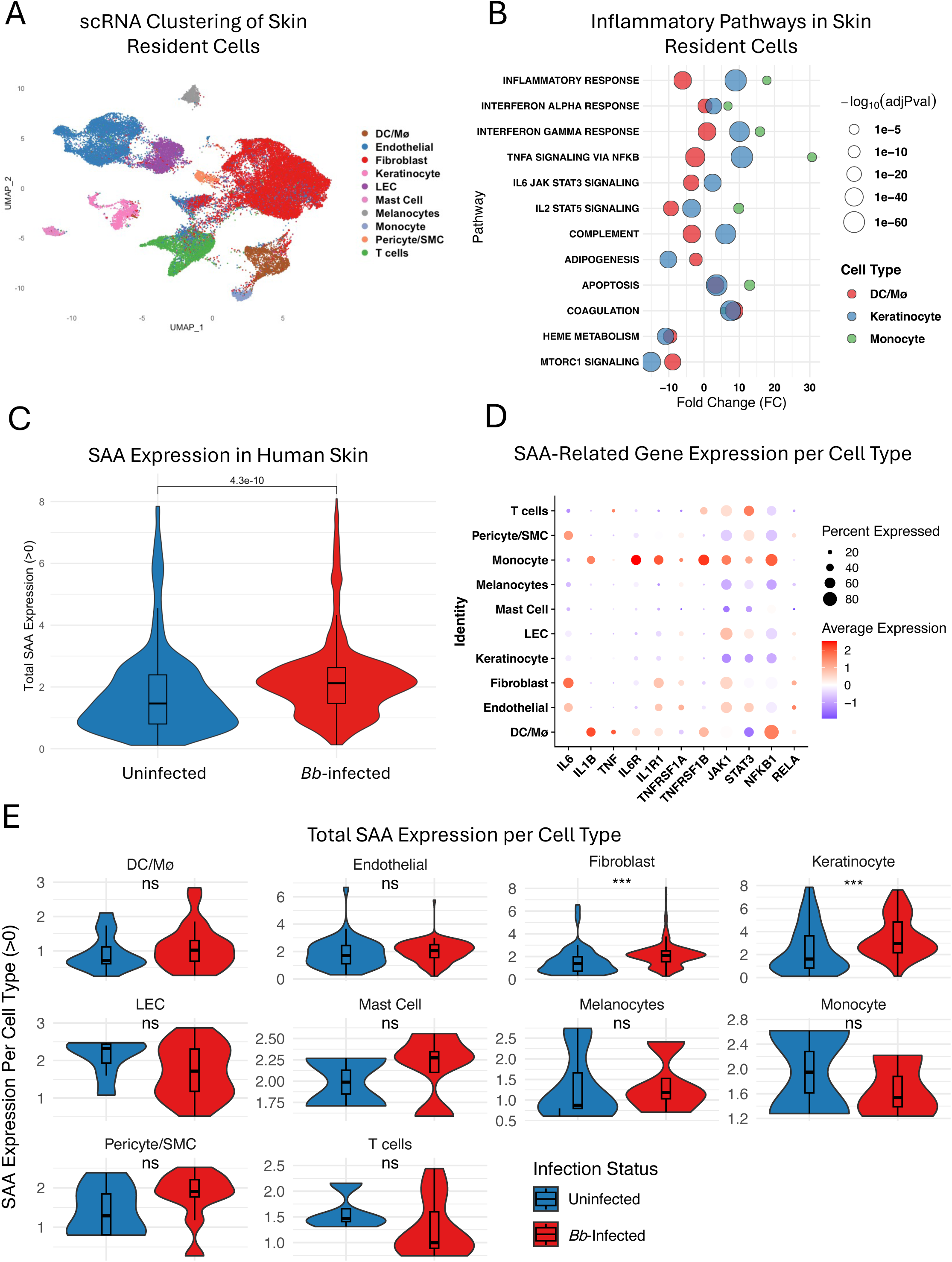
Hub Gene Analysis of Adipose Tissue Adipose Stem/Progenitor Cells (ASPCs) (Cluster 5) and Lymphatic Endothelial Cells (LECs) (Cluster 10). (A) Network visualization showing genes as nodes (size corresponds to degree; color indicates log₂FC: red = upregulated, blue = downregulated in infection). Connecting lines represent predicted functional associations.

**Supplementary Figure S5.**
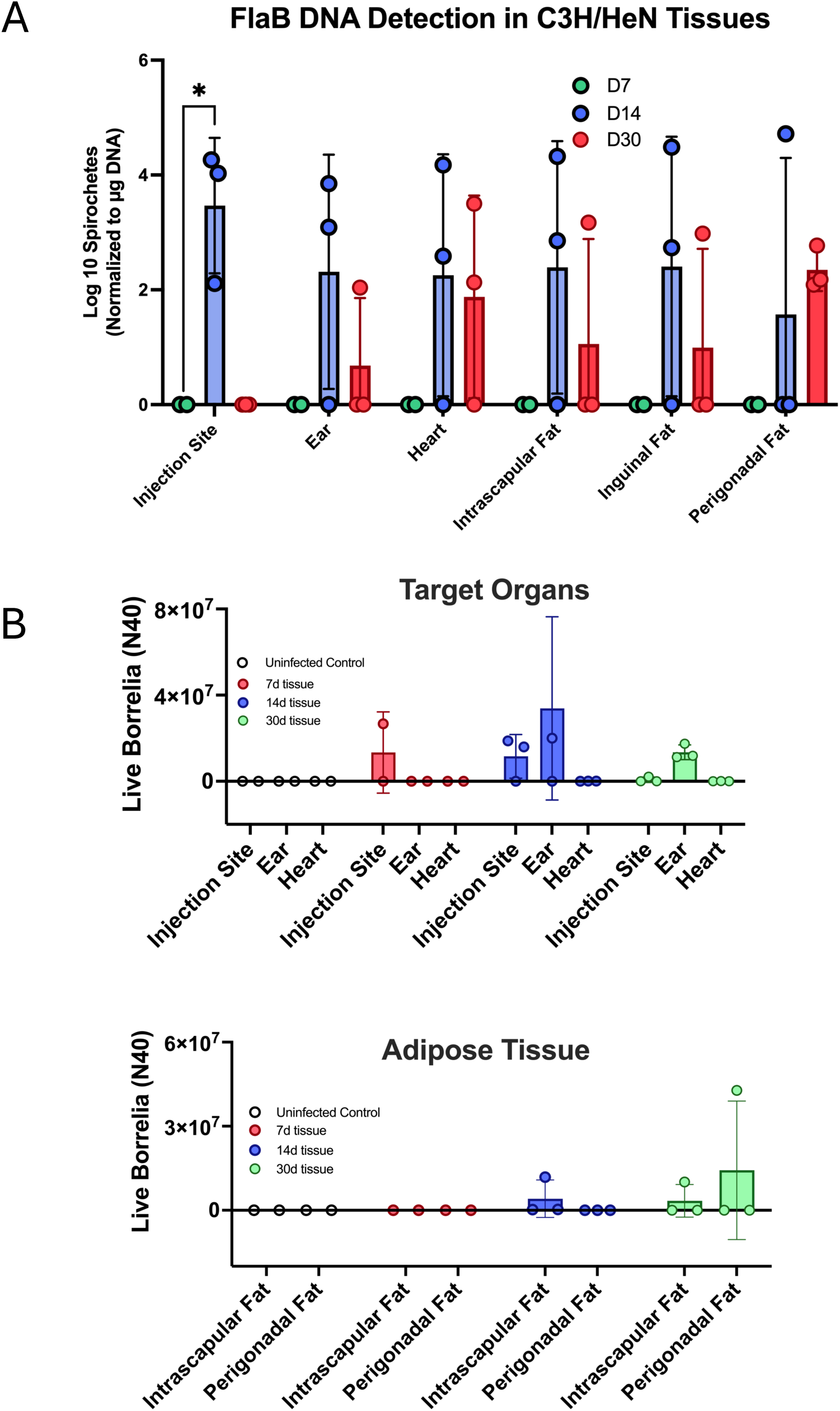
*Borrelia burgdorferi* N40 infection in C3H/HeN mice. Mice were infected with 10^5^ live spirochetes and tissues were harvested at 7d, 14d, and 30d post infection. n=3 per timepoint. Figures show (A) quantitative PCR loads detecting DNA copies of *Borrelia* flagellin (*FlaB*) sequences from target site and adipose depots. (B) Live *Borrelia* cultures from small portions of tissue harvested from mice at every timepoint were incubated in clean BSKII media supplemented with fosfomycin, rifampicin, and amphotericin B to prevent contamination. Cultures were incubated for 14d and checked for live spirochetes via dark field microscopy at 40x magnification using a Neubauer Improved hemocytometer.

**Supplementary Figure S6.**
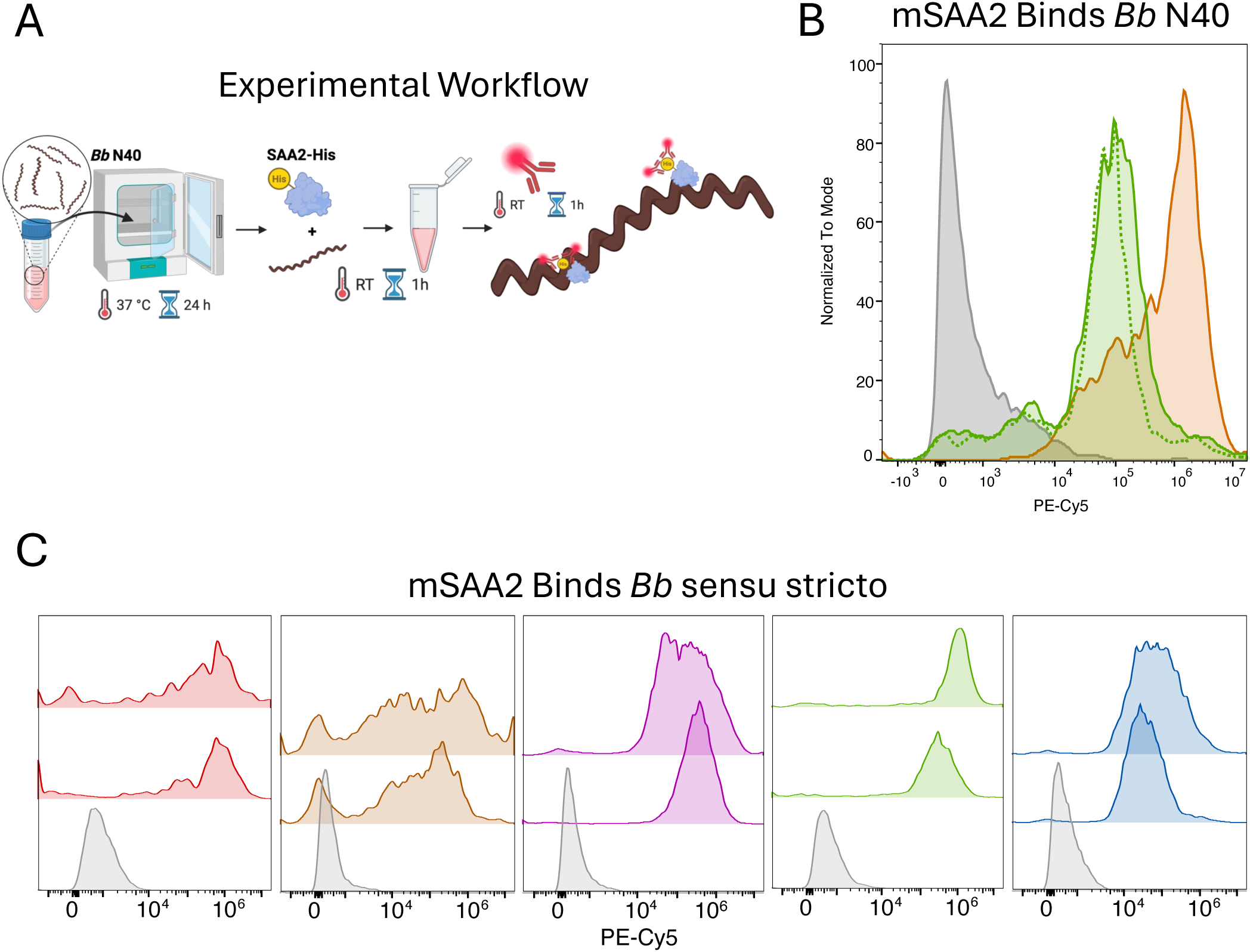
Murine SAA2 Binds *Borrelia burgdorferi ss.* (A) Experimental workflow diagram of flow cytometry binding assay. Spirochetes cultured at 37°C for 24h were allowed to bind recombinant His-tagged mSAA2. Anti-His tag fluorescent (PE/Cy5) primary conjugated antibodies were bound to SAA-spirochete complexes to label protein attached to spirochetes. Spirochetes were washed and used for flow cytometry analysis compared to unstained spirochete complexes. (B) Histogram plots indicating murine SAA2 (mSAA2) binds SAA2 relative to staining control (*Bb* + mSAA2) and positive control protein peptidoglycan recognition protein-1 (PGLYRP1 ref 39). (C) Binding of mSAA2 to *Bb* sensu stricto strains B31, CA8, HP19, CT-1, NT-1.

**Supplementary Figure S7.**
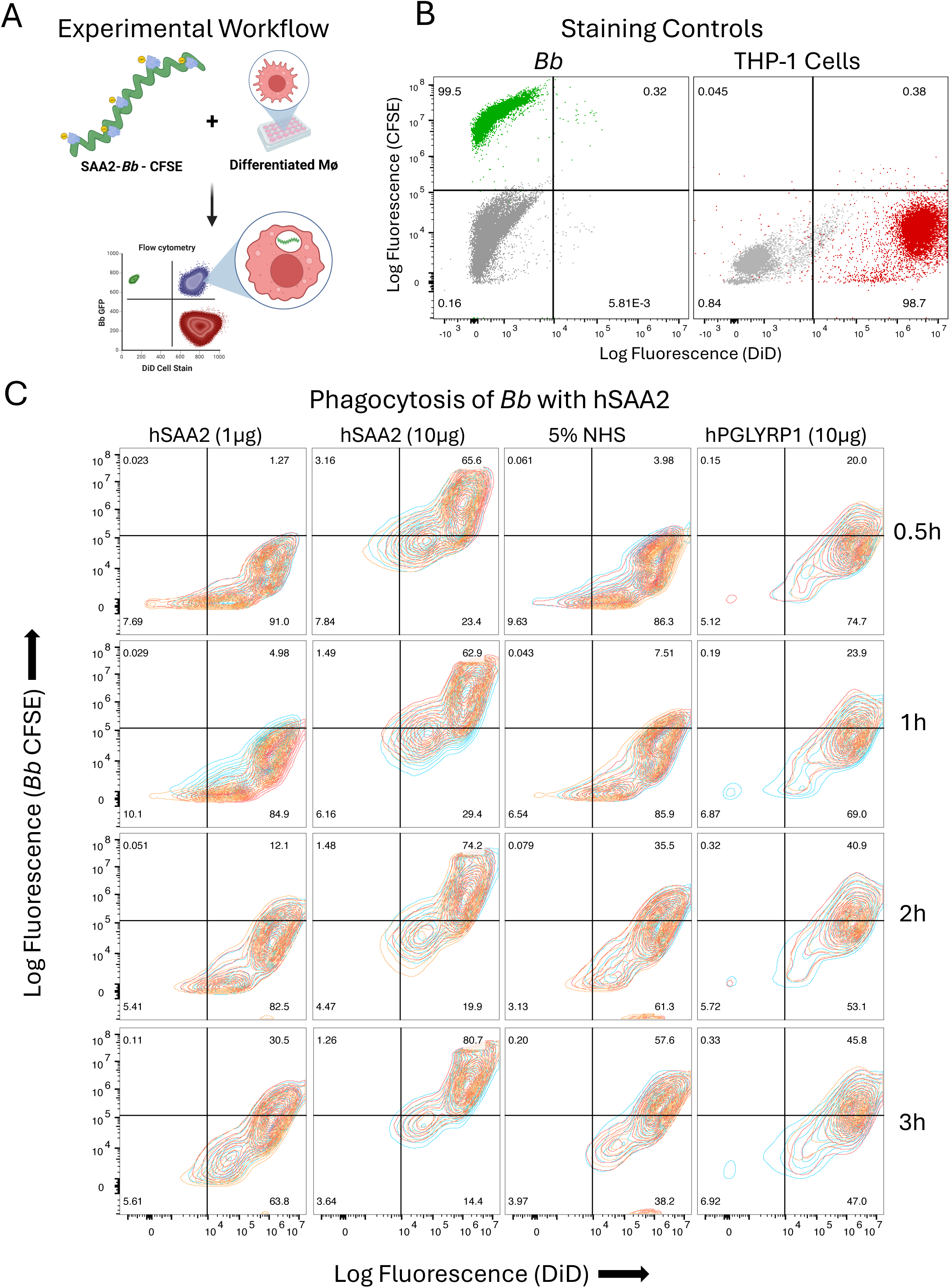
Flow Cytometry Plots Related to Figure 3. (A) Workflow diagram of labeled spirochetes added to stained cells to be measured by flowcytometry. Spirochetes were allowed to bind SAA2 previously described (Fig 1) and labeled with CFSE. Separately, differentiated THP-1 macrophage-like cells were labeled with membrane stain DiD. Labeled spirochetes were added to stained cells and harvested at 0.5-3.0h post incubation at 37°C. (B) Flow cytometry controls of CFSE-stained *Bb* N40 spirochetes (green) and DiD-labeled cells (red) showing detection capability of both positive- and negative-stained populations. (C) Contour plots of THP-1 cells and spirochetes incubated with either hSAA2, 5% normal human serum (NHS), and hPGLYRP1 (10µg) comparing levels of double positive populations over time (0.5, 1.0, 2.0, 3.0h in co-culture). Contour plots show percentages of cells in each quadrant in which triplicate plots overlayed. Each replicate value from quadrant 2 (top right) used for quantification (Figure 3A).

**Supplementary Figure S8.**
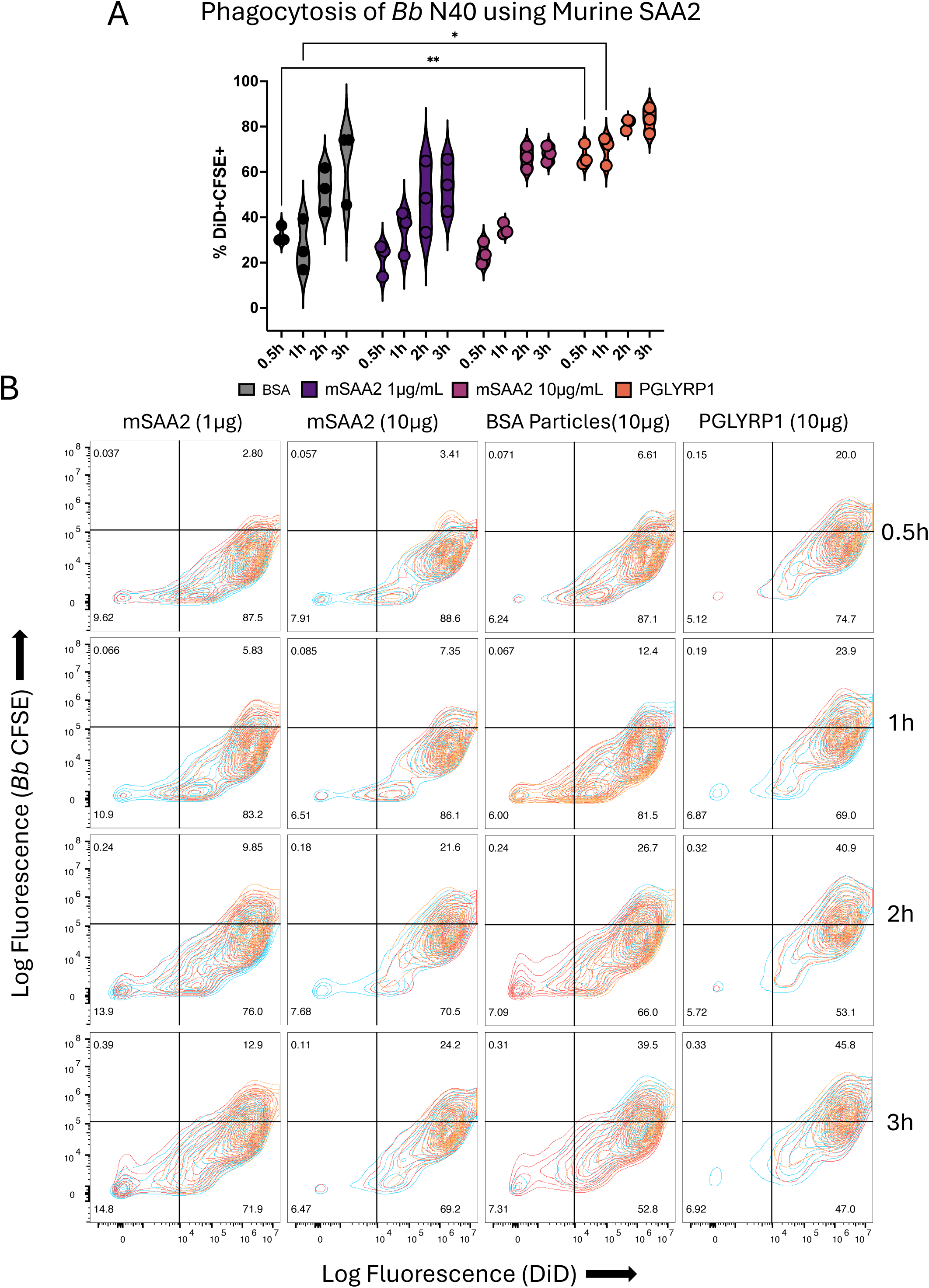
Phagocytosis of *Bb* N40 Using mSAA2. (A) Violin plot generated of the proportion of double positive (DiD+CFSE+) populations after co-incubating mSAA2-labeled spirochetes with differentiated THP-1 macrophages for 0.5h, 1h, 2h, and 3h, as performed in Supplementary Figure S2 A-B. Statistical significance was determined a two-way ANOVA followed by post hoc Tukey test to correct for multiple comparisons. (B) Flow cytometry contour plots of BSA particles (in 1x PBS) (10µg), hPGLYRP1 (10µg) positive control, and mSAA2 (1µg or 10µg) treated spirochetes incubated with differentiated THP-1 macrophages for 0.5h, 1h, 2h, 3h co-culture. Contour plots show percentages of total cells in each quadrant in triplicate. Values from quadrant 2 (top right) used for quantification (A).

